# How do noise and precision contraints jointly determine the minimum sketch size for fixed-size MinHash Jaccard estimation?

**DOI:** 10.64898/2026.08.15.744971

**Authors:** Anicet E. T. Ebou, Dominique K. Koua

## Abstract

MinHash-based genome comparison is governed by two statistical constraints: a noise floor on *k*-mer size, controlling chance collisions between unrelated sequences, and a precision floor on sketch size, controlling uncertainty in the estimated Jaccard similarity. These constraints are best characterized for fixed-size bottom-sketch MinHash, the classical construction underlying tools such as Mash and still the standard basis for comparing genomes of similar size. Nevertheless, while, the noise floor has an established closed-form solution, sketch size is commonly selected using fixed defaults or heuristic choices independent of genome length and target precision. Moreover, recent work on scaled (FracMinHash) sketching has highlighted the difficulty of obtaining an analogous closed-form confidence interval for the Jaccard similarity. Here, we show that under the fixed-size bottom-sketch MinHash model, the shared-hash count follows a binomial model, which lets classical proportion-estimation theory be applied directly. Combining this with Fofanov’s *k*-selection criterion and the exact identity-Jaccard relationship yields a single closed-form design rule, *s*\*(*L, ρ*\*, *α*), that unifies the noise and precision floors into one operating envelope. The resulting design rule was validated against exact Clopper-Pearson interval inversion, idealized binomial sampling, realistic overlapping *k*-mer simulation and 54 bacterial genome pairs spanning species-to-order taxonomic levels. In realistic simulations, empirical coverage remained within 1-3 percentage points of the idealized reference in 14 of 15 tested cells. In real genomes, the classical identity-Jaccard relationship showed increasing positive bias with taxonomic divergence, from a median of −1.4% within species to +27% at genus and +77% at family level. We further show that *s*\*(*L*) is not smooth in genome length but a discontinuous, previously unreported staircase caused by the ceiling function used for *k*-mer selection, with practical consequences concentrated at specific genome-size boundaries.

## 1 Introduction

Exact *k*-mer-based comparison scales poorly as sequence collections grow, since all-pairs comparison requires *O* (*N* ^2^) operations on potentially large *k*-mer sets [Steinegger and Söding, 2018]. MinHash [Broder, 1997] addresses this by retaining only the *s* smallest hash values from a set’s fingerprint, so that the Jaccard index between two sets can be estimated from a fixed-size sample rather than the full intersection. Ondov et al. [2016] adapted this bottom-sketch construction to *k*-mer sets in Mash, coupling the Jaccard estimate to a Poisson mutation model to report a genome-wide distance, and demonstrating all-pairs comparison of the 54,118 RefSeq genomes in under 33 CPU-hours. This combination of efficiency and biological interpretability drove wide adoption of sketch-based comparison (Mash, sourmash [Pierce et al., 2019], Dashing [Baker and Langmead, 2019], BinDash [Zhao, 2019]).

Two independent statistical constraints govern the validity of such a sketch, developed along separate tracks for two decades. The first is a constraint on *k*: it must be large enough that two unrelated sequences share almost no *k*-mers by chance. Fofanov et al. [2004] derived this directly, showing that for a genome of length *L* the expected number of chance-shared *k*-mers falls below a fixed tolerance once *k* exceeds *k*\*(*L*) = ⌈log_4_(100*L*)⌉, a criterion that predates MinHash-based comparison and has since been invoked broadly, including in *k*-mer-based population-genetic and pangenomic methods [Roberts et al., 2025] and biodiversity genomics [Jenike et al., 2025]. This criterion controls chance *k*-mer matches but does not determine how many *k*-mers must be *sampled* into a sketch for the resulting Jaccard estimate to be statistically informative. Indeed, *k* and *s* govern different sources of uncertainty, yet are practically linked, since *k* sets the expected similarity *J* while *s* governs the sampling uncertainty of its estimator.

The second constraint concerns this sampling question directly, and has received comparatively little explicit treatment in genomics despite being a classical problem in survey sampling, where the sample size required to estimate a proportion at a target relative precision follows directly from Cochran [1977]. This inversion is not itself a new statistical result, our contribution is to compose it with Fofanov’s criterion and the identity-Jaccard relationship into a single genome-length- and divergence-aware design formula, and to characterize the consequences of doing so. Ondov et al. [2016] note that the Jaccard estimate’s error bound depends only on *s*, independent of genome length, and supply a significance test for chance *k*-mer sharing, but sketch size itself is left a heuristic (with a suggested *s* = 1000 for bacterial genomes) disconnected from any target precision, genome length, or resolvable divergence, and driving other tools choice.

Later, MinHash extensions addressed a different limitation: poor behaviour when comparing sets of very different cardinality. Koslicki and Zabeti [2019] introduced containment MinHash, and Irber [2020], Irber et al. [2022] described FracMinHash, in which a scale factor rather than a fixed sketch size determines which hashes are retained (independently described as the *universe minimizer* [Ekim et al., 2021] and *mincode syncmers* [Edgar, 2021]). This formulation enabled statistical treatment of containment estimates. Building on the *k*-mer mutation statistics of Blanca et al. [2022], Hera et al. [2023] derived an unbiased FracMinHash containment estimator with a closed-form confidence interval, but report that the analogous derivation for the Jaccard index fails to establish asymptotic normality, leaving no closed-form Jaccard interval under the FracMinHash construction (Hera et al. 2023, Supplemental A.7). This leaves an asymmetry in the literature: precision-aware interval estimation exists for containment under variable-size sketching, but not for Jaccard under either variable- or fixed-size sketching, despite the latter remaining widely used. The noise floor and precision floor have not, to our knowledge, been expressed as a single operating envelope for sketch design.

In this study we unify these two constraints for fixed-size bottom-sketch MinHash, where the shared-hash count is binomially distributed given *s* and *J* and no ratio of dependent random denominators arises. Composing classical sample-size theory with Fofanov’s criterion and the identity-Jaccard relationship [Blanca et al., 2022] yields a closed-form formula, *s*\*(*L, f*_min_, *ρ*\*, *α*): given a genome length *L*, target relative precision *ρ*\*, confidence level 1 − *α*, and *k* selected by Fofanov’s criterion, what is the minimum sketch size satisfying the resulting precision constraint? We validate this formula against exact Clopper-Pearson inversion and realistic sketching simulation, and show two previously unreported consequences of the unification: a structural one, that *s*\*(*L*) is discontinuous in genome length rather than smooth, arising directly from the ceiling function in *k*-mer selection; and a biological one, that the identity-Jaccard relationship departs substantially from real bacterial genome behaviour below the species level, in a pattern structured by taxonomic rank.

## 2 Materials and Methods

### 2.1 Statistical framework

Under the bottom-sketch construction, the shared-hash count *x* is hypergeometrically distributed, *x* ~ Hypergeometric(*N, K, s*) with *N =* |*A ∪ B* |, |*A ∩ B*|, since the *s* retained ranks are drawn without replacement from this finite population [Blanca et al., 2022]. We model *x* as Binomial(*s, J*) with *J* = *K/N* throughout, the standard limiting approximation as *N*→ ∞. Because hypergeometric variance *sJ* (1 − *J*)(*N* − *s*)*/*(*N* − 1) is always ≤ binomial variance at the same *N, K, s*, this makes the resulting interval conservative rather than anti-conservative, and negligibly so given *N* ≫ *s* throughout this work (Supplementary S1 and S10).

We compute the exact Clopper-Pearson confidence interval on *Ĵ* = *x/s* as

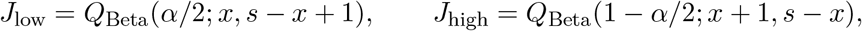

and define *ρ* = (*J*_high_ − *J*_low_)*/*(2*Ĵ*). For *x* = 0 or *x* = *s, J*_low_ = 0 or *J*_high_ = 1 respectively, so coverage estimates remain unconditional and comparable to nominal 1 −*α*.

The Wald approximation 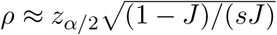 [Cochran, 1977] inverts to

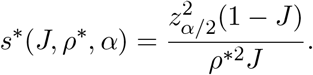

Under the simple mutation model [Blanca et al., 2022], each nucleotide is independently substituted with probability *r*, giving *q* = 1 − (1 − *r*)^*k*^ and unsketched Jaccard similarity (1− *q*)*/*(1 + *q*) exactly. Defining *f*_min_ := 1− *r*, the minimum per-site identity between the genomes compared, this yields the exact identity-Jaccard relationship

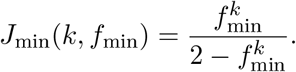

*f*_min_ is the user-specified design threshold throughout, except in the real genome benchmark (Section 2.7), where it is instantiated retrospectively as each pair’s FastANI-estimated 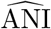 Substituting *k* = *k*\*(*L*) = ⌈log_4_(100*L*) ⌉ [Fofanov et al., 2004] and *J* = *J*_min_(*k*\*(*L*), *f*_min_) gives the joint closed form used throughout. The resulting *s*\* is conditional on the validity of this identity-Jaccard relationship.

### 2.2 MinHash sketch construction

Let *H*(*A*), *H*(*B*) denote the complete hashed *k*-mer sets of two genomes and bottom_*s*_(·) the *s* smallest elements under hash order. We compute *H*_*A*_ = bottom_*s*_(*H*(*A*)), *H*_*B*_ = bottom_*s*_(*H*(*B*)), form *H*_*AB*_ = bottom_*s*_(HA ∪ HB), and take *x* = |*HAB ∩ HA ∩ HB*|. This is exactly equivalent to bottom_*s*_(*H*(*A*) ∪ *H*(*B*)), the true bottom-*s* sketch of the full union (Supplementary S11), matching standard practice in Mash and sourmash.

Each *k*-mer hash lies in {0, …, 2^62^ − 1} : two independently seeded 32-bit MurmurHash values are masked to {0, …, 2^31^ − 1} and combined as *h* = *h*_high_ 2^31^ + *h*_low_. No hash collisions were observed in any analyzed dataset, consistent with the 62-bit space vastly exceeding per-genome *k*-mer counts.

Because a sketch size must be a non-negative integer, we use

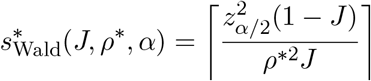

in every simulation and real-data analysis (Section 2.4 - Section 2.7), the unrounded form is used only in Section 2.3 for direct comparison against 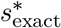. When the available hash union is smaller than *s*\*, the full available set is used instead, giving effective size *s*_used_ = min(*s*\*, |*H*_*A*_ ∪ *H*_*B*_|). We define a comparison as *degenerate* when *x* = 0 or *x* = *s*_used_ (*ρ* undefined), reported as a distinct category rather than assigned a reliability tier.

### 2.3 Wald approximation versus exact interval inversion

For candidate sketch size *s*, we define the deterministic exact relative half-width

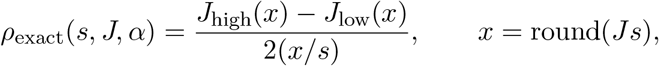

using the plug-in expected count rather than a random draw, and

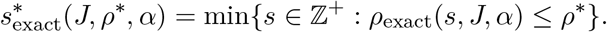

Because *x* = round(*J s*) introduces integer rounding, *ρ*_exact_ is not guaranteed monotonic in *s*. Consequently, we identify 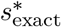 via an exhaustive vectorized scan (adaptively bounded, up to *s*_max_ = 5 × 10^6^) rather than an algorithm assuming monotonicity, evaluated in parallel across CPU cores (Supplementary S2). Of 360 grid cells, 19 (5.3%) were infeasible within *s*_max_ and excluded. We separately report 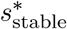 the smallest *s* beyond which the target holds for all larger *s* within range, to quantify how often non-monotonicity causes 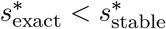.

### 2.4 Idealized sketching calibration

To confirm 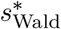 under the idealized assumption (*x* ~ Binomial(*s*\*, *J*) exactly), we drew *n* = 10,000 samples per cell across *L* ∈ [10^5^, 10^10^] (30 log-spaced points) and *f*_min_ ∈ {0.8, 0.9, 0.95, 0.99, 0.999} at *ρ*\* = 0.10, *α* = 0.05, recording coverage and realized *ρ* (including degenerate draws). A secondary sweep evaluated *ρ*\* ∈ {0.05, 0.10, 0.30}, *α* ∈ {0.01, 0.05, 0.10} at *L* = 1 Mb, *f*_min_ = 0.9.

### 2.5 Realistic-sequence sketching calibration

We simulated genome pairs under the simple point-mutation model [Blanca et al., 2022, Hera et al., 2023]: for each cell (*L* ∈ {10^4^, 10^5^, 10^6^}, *f*_min_ ∈ {0.8, 0.9, 0.95, 0.99, 0.999}), an i.i.d. reference sequence was generated and mutated at each position independently with probability *p* = 1 − *f*_min_ (uniform over the three alternative bases, no indels). For each replicate (*n* = 10,000/cell), *k*-mers were hashed as in Section 2.2 at *k* = *k*\*(*L*), and compared via the bottom-sketch procedure. We recorded, per replicate: the exact realized *J*_exact_ from the full unsketched *k*-mer sets; coverage/*ρ* of *Ĵ*against *J*_exact_; and coverage/*ρ* against the fixed prediction *J*_theory_ = *J*_min_(*k*\*(*L*), *f*_min_), alongside a matched idealized Binomial (*s*\*, *J*_theory_) reference.

### 2.6 Staircase structure of *s*\*(*L*)

*s*\* was evaluated on 2,000 log-spaced genome lengths spanning *L* ∈ [10^5^, 10^10^] per *f*_min_ at *ρ*\* = 0.10, *α* = 0.05. Step edges (consecutive grid points differing in *k*\*(*L*)) were checked against the theoretical boundary *L* = 4^*k*^ */*100, and step ratios 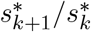 against the small-*J* prediction 1*/f*_min_ (Section 2.1).

To assess how frequently real genomes fall near a boundary, we classified every genome in the complete GTDB release 232 census (901,341 genomes) by proximity of its reported genome size to the nearest *L* = 4^*k*^ */*100, repeated for the full set and for species representatives alone (Supplementary S6).

### 2.7 Real genome benchmark

Genome pairs spanning a wide ANI range were sampled from GTDB R232 [Parks et al., 2022] via xgt v1.1 [Ebou et al., 2026], following a taxonomically anchored design adapted from Hera et al. [2023]. Specifically, for eight anchor bacterial species, the GTDB representative genome was retrieved and three further genomes sampled at each of species, genus, family, and order rank, selected to share that rank with the anchor but not the next more specific rank (Table S1). Sequences were downloaded via `datasets` v18.34.0 [Cox et al., 2025], and 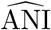 was obtained via FastANI v1.33 [Jain et al., 2018], with 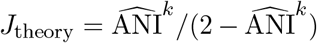, excluding pairs below FastANI’s ~ 75–80% resolution floor [Hera et al., 2023]. For each pair, *k* was selected via *L* = max (*L*_*A*_, *L*_*B*_), a conservative choice given Fofanov’s criterion’s natural two-genome generalization depends on *L*_*A*_*L*_*B*_ rather than max (*L*_*A*_, *L*_*B*_)^2^ (Supplementary S7 and S12). The two selections coincided for all 54 retained pairs.

Assembly completeness and plasmid presence (via header string matching) were recorded but not used for exclusion (assembly statistics reported in Results). This benchmark is retrospective as 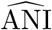 is used as the plug-in *f*_min_. Therefore, results characterize how *s*\* behaves relative to a pair’s own realized similarity rather than prospective design performance (Section 2.8 addresses the latter directly). Comparisons were performed at *s* = 1000 and at formula-recommended *s*\* (*ρ*\* = 0.10, *α* = 0.05), recording *ρ*, coverage of *J*_exact_, and reliability tier (reliable *ρ* ≤ 0.10; borderline ≤ 0.30; unreliable *>* 0.30; degenerate as defined in Section 2.2). Each distinct (accession, *k*) combination was hashed once and cached, since genomes recur across pairs.

### 2.8 Prospective validation at fixed design targets

To test performance without prior knowledge of a pair’s true divergence, we repeated the comparison using two fixed targets, *f*_min_ ∈ {0.90, 0.95}, applied identically to all 54 pairs regardless of 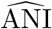. *k* was unchanged from Section 3.6; 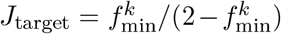 and 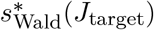 were used to construct the comparison from the same cached hash sets, evaluated against each pair’s *J*_exact_. Pairs were stratified by the gap between 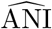 and the fixed target (at/above target; up to 5pp below; more than 5pp below).

### 2.9 Software

Analyses were implemented in R v4.6.1 [R Core Team, 2025] using `qbeta/qnorm, furrr` v0.4.0/ `future` v1.75.0 for parallel Monte Carlo replication, and `ggplot2` v4.0.3 [Wickham, 2016] for visualization.

## 3 Results

### 3.1 Closed-form sketch-size formula

Composing classical proportion-estimation theory with Fofanov’s *k*-selection criterion and the identity-Jaccard relationship yields the closed-form design rule used throughout:

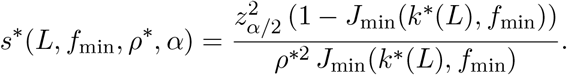

The derivation was independently re-verified step by step and validated numerically against exact Clopper-Pearson inversion below (Supplementary S1). The small-*J* approximation used in the staircase analysis (Section 3.5), *J*_min_(*k, f*) ≈ *f* ^*k*^*/*2, has relative error exactly equal to *J*_min_(*k, f*) itself, accurate to within a few percent at high divergence but directional only near sequence identity (Supplementary S1).

### 3.2 Wald approximation versus exact interval inversion

Across the full evaluated grid (341 of 360 cells valid), 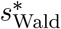 tracked the true exact minimum 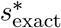 closely and, with a single exception attributable to integer-rounding non-monotonicity rather than any deficiency of the approximation, conservatively. Specifically, median relative error was 0.8-3.8% at the precision targets most relevant to genome comparison (*ρ*\* ≤ 0.10), rising to 109.6% only at the loosest setting tested (*ρ*\* = 0.30, *α* = 0.10), where absolute sketch sizes are correspondingly small and the discrepancy is of limited practical consequence (Table S2). We adopt 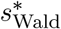 as the recommended design formula throughout, with the exact inversion retained solely as a validation reference (Supplementary S2).

### 3.3 Calibration under the Binomial model

Under the idealized sampling assumption, 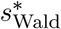 delivered coverage consistent with nominal across the full grid of genome lengths and minimum identities at *ρ*\* = 0.10, *α* = 0.05. Coverage ranged from 0.9475 to 0.9984 (median 0.9553; 150 cells), with no cell falling below nominal by more than three Monte Carlo standard errors, and median realized *ρ* = 0.1017 (Table S3, Figures S1-S2). A secondary sweep across (*ρ*\*, *α*) at *L* = 1 Mb, *f*_min_ = 0.9 showed the same pattern, with median *ρ* consistently at or slightly above target. Median overshoot 1.65% across the full grid and went up to 145.8% only at the smallest tested sketch sizes, *s*\* ∈ {10, 11, 12}, where the discreteness of achievable *ρ* values is severe enough that no integer outcome lands near the target, even though coverage itself remains intact, 0.978-0.998, in these same cells (Supplementary S3). Correspondingly, the proportion of individual replicates achieving *ρ* ≤ *ρ*\* was below 50% in every cell, a discreteness artifact rather than a calibration concern (Supplementary S3). These results confirm 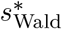 is well calibrated under the model’s own idealized assumptions across five orders of magnitude in genome length and the full range of minimum identities considered.

### 3.4 Calibration on simulated genome sequences

Coverage against each replicate’s exact realized Jaccard index *J*_exact_ closely matched the idealized binomial reference across the full simulation grid (*L* ∈ {10^4^, 10^5^, 10^6^}, *f*_min_ ∈ {0.8, …, 0.999}). More precisely, only 1 of 15 cells exceeded a 2pp coverage gap (and in the conservative direction), and median realized ρ matched the idealized value to within 4% everywhere (Table 1).

**Table 1.** Coverage against the fixed formula prediction (*J*_theory_), each replicate’s exact realized Jaccard index (*J*_exact_), and the idealized binomial reference, averaged across the three tested genome lengths at each minimum identity. Target coverage is 0.95.

| $f_{\min}$ | vs. $J_{\text{theory}}$ | vs. $J_{\text{exact}}$ | vs. idealized |
| --- | --- | --- | --- |
| 0.999 | 0.987 | 0.990 | 0.989 |
| 0.99 | 0.956 | 0.964 | 0.959 |
| 0.95 | 0.933 | 0.956 | 0.955 |
| 0.90 | 0.909 | 0.956 | 0.953 |
| 0.80 | 0.630 | 0.967 | 0.954 |

By contrast, coverage against the fixed formula prediction *J*_theory_ degraded substantially at higher divergence (0.59-0.66 at *f*_min_ = 0.8), fully explained by a corresponding, monotonically growing divergence between *J*_theory_ and mean realized *J*_exact_ (0.005-0.012% at *f*_min_ = 0.999 to 8.0-8.1% at *f*_min_ = 0.8; Table S4). This result reflect a known limitation of the simple point-mutation model at higher divergence and is consistent with the overestimation of mutation distance by the classical Mash model reported independently by Hera et al. [2023]. This bias is favorable for practical use. As realized *J*_exact_ runs consistently above *J*_theory_ at high divergence, median realized *ρ* remains at or below target throughout the tested range, so the formula does not underprovision precision even where its identity-Jaccard prediction is imprecise. *s*\* therefore delivers its target precision guarantee reliably on real sketched sequences *when the assumed design target is met or exceeded by the pair being compared* (Section 3.7). For pairs whose true divergence exceeds the assumed target, coverage can fall materially below nominal, a limitation distinct from, and additive to, the identity-Jaccard bias characterized in Section 3.6.

### 3.5 Staircase structure of the minimum sketch size

Because *k*\*(*L*) = ⌈ log_4_(100*L*) ⌉ is a ceiling function, *s*\*(*L*) is piecewise constant within each Fofanov *k*-band and discontinuous at band edges, confirmed directly on a dense grid (*L* ∈ [10^5^, 10^10^]). Specifically, *s*\* remained exactly flat within each band and jumped discretely at every one of 40 detected transitions across the five minimum identities tested (Figure 1, Figure S4).

**Figure 1.**
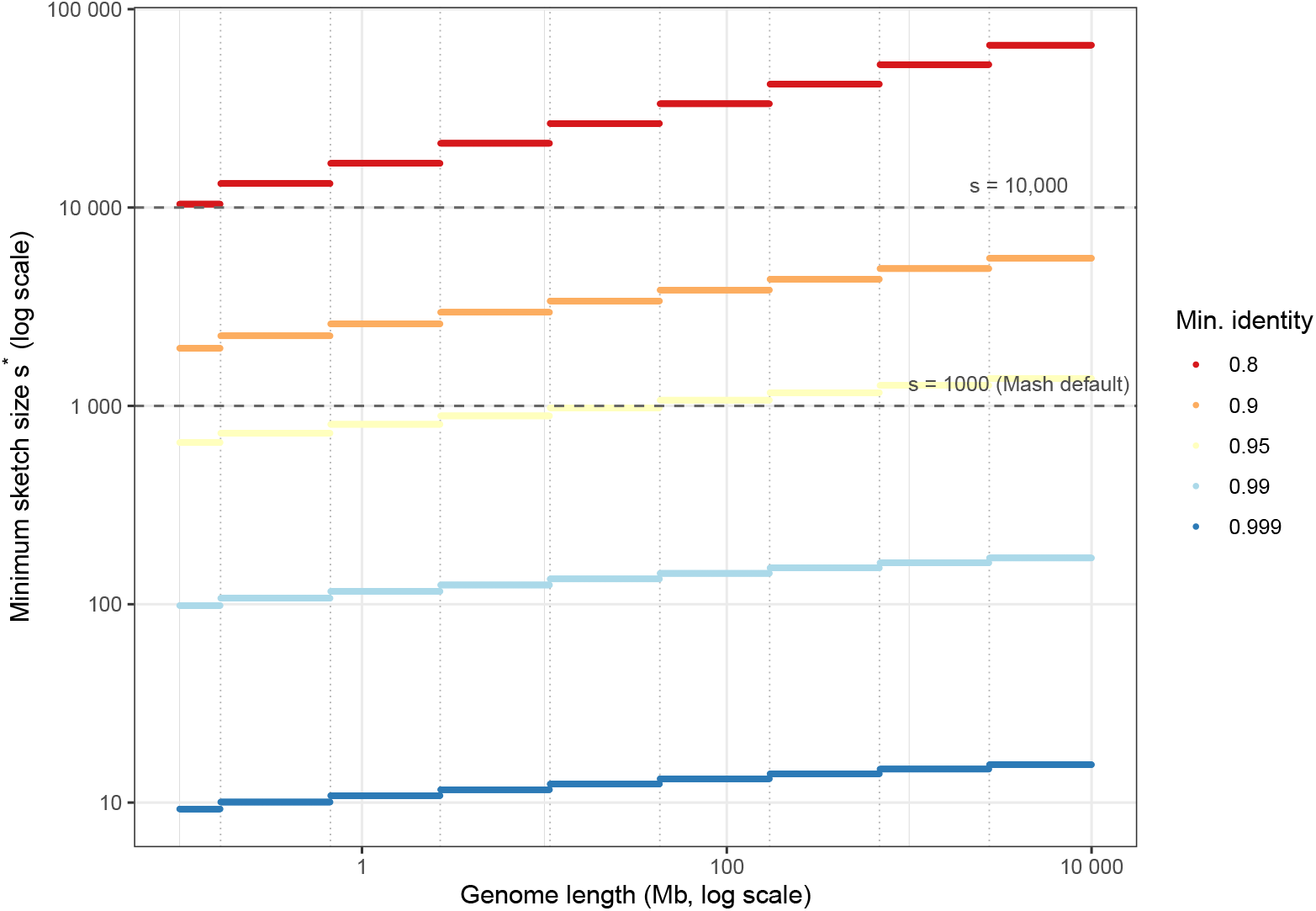
Minimum sketch size *s*\* vs. genome length *L*, at *ρ*\* = 0.10, *α* = 0.05, five minimum identities *f*_min_. *s*\* is piecewise constant within each *k*-band and jumps at band edges (dotted lines, *L* = 4^*k*^*/*100); points, not lines, are plotted to avoid implying interpolation across a discontinuity. Dashed lines mark *s* = 1000 and *s* = 10,000 for reference.

The closed-form boundary *L* = 4^*k*^ */*100 fell inside the grid bracket for all 40 transitions (max bracket width 0.58% of *L*_predicted_; Table S5), confirming it as exact rather than approximate. Observed step-height ratios 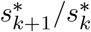 agreed with the small-*J* prediction 1*/f*_min_, improving with divergence as expected (median relative error 0.65% at *f*_min_ = 0.8 rising to 6.41% at *f*_min_ = 0.999; Table 2).

**Table 2.** Step-height ratio agreement across all 8 transitions per minimum identity.

| $f_{\min}$ | Median rel. error (%) | Max rel. error (%) |
| --- | --- | --- |
| 0.8 | 0.65 | 1.48 |
| 0.9 | 2.43 | 3.94 |
| 0.95 | 4.12 | 5.88 |
| 0.99 | 5.94 | 7.80 |
| 0.999 | 6.41 | 8.28 |

This structure is a direct, and in retrospect elementary, consequence of *k*\*(*L*) being a ceiling function. Nonetheless, we are not aware of it being stated explicitly in the MinHash sketch-size literature.

Against the complete GTDB genome census, 99.06% of genomes fell within a single *k*\* band and were structurally unaffected, but 7.6% sat within 5% of a boundary and 15.5% within 10%, essentially unchanged when restricted to species representatives alone (7.8%, 15.5%; Supplementary S6).

### 3.6 Evaluation on real bacterial genomes

As a retrospective calibration check-using each pair’s 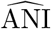 as the plug-in *f*_min_, 54 genome pairs spanning 77.6% to 99.9% ANI were sampled from GTDB across eight taxonomically anchored bacterial lineages, comparing reliability-tier classification at *s* = 1000 against the formula-recommended *s*\*.

The two sketch sizes produced markedly different classifications: 47 of 54 pairs (87.0%) were reclassified (Table 3). At *s* = 1000, 13 pairs (24.1%) were unreliable and at *s*\*, none were, with the four smallest-*s*\* pairs 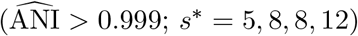 instead degenerate. In each of these degenerate cases *s*_used_ = *s*\* exactly and *x* = *s*_used_ (complete saturation), leaving *ρ* undefined rather than indicating unreliability. The proportion classified reliable increased from 53.7% to 59.3%. This reclassification should not, however, be read as evidence that *s*\* precisely targets the sketch size each pair required.

**Table 3.** Reliability-tier classification of 54 real GTDB genome pairs, *s* = 1000 vs. formula-recommended *s*\*. No comparisons were degenerate at *s* = 1000.

| Tier | $s = 1000$ | $s^*$ |
| --- | --- | --- |
| Reliable | 29 | 32 |
| Borderline | 12 | 18 |
| Unreliable | 13 | 0 |
| Unreliable (degenerate) | 0 | 4 |
| **Total** | **54** | **54** |

Comparing *J*_theory_ against *J*_exact_ revealed a bias far larger than on simulated sequences (Figure 2): median +7.9% (range −12.9% to +275%), with 18.5% of pairs showing *J*_exact_ more than double *J*_theory_. Unlike the smooth, modest simulated-sequence bias (≤ 8%), this was strongly structured by taxonomic rank: species-level pairs (*n* = 24) showed 1.4% median bias (median 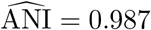), closely matching simulation, while genus-level (*n* = 17) showed +27.1% 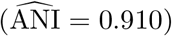 and family-level (*n* = 13) showed +77.1% 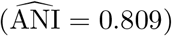. All pairs exceeding 100% bias were genus- or family-level (Table S7). The pattern held within individual anchor lineages (e.g. *E. coli*: −4.2% → +18.6% → +66.4% across the same three ranks).

**Figure 2.**
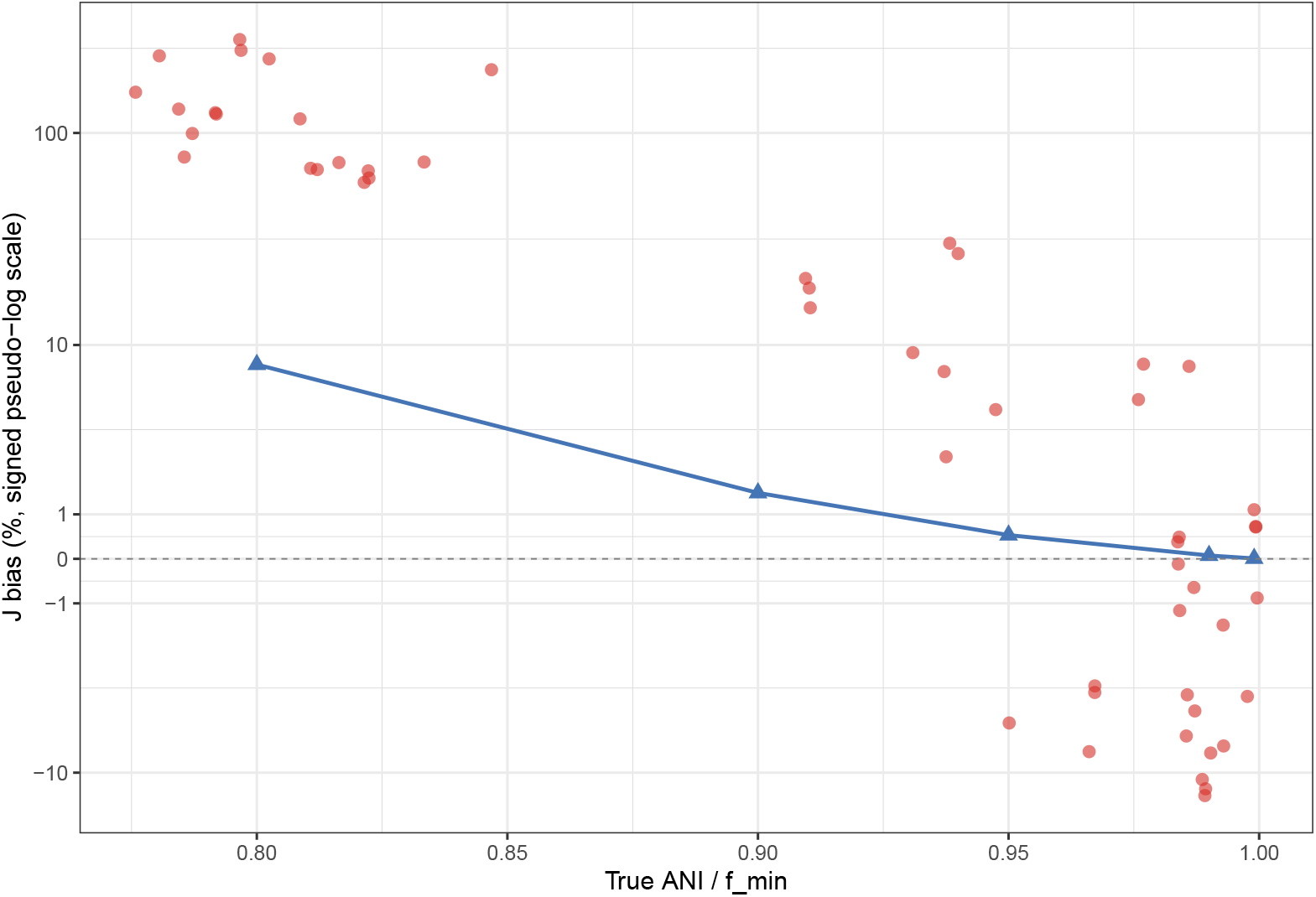
Bias of *J*_theory_ against *J*_exact_ vs. minimum identity. Blue: simulated sequences (Section 3.4), bias ≤ 8%. Red: 54 real GTDB pairs (Section 3.6), departing sharply below the species boundary, exceeding 200% at genus/family level.

### 3.7 Prospective validation at fixed design targets

The benchmark above is retrospective and cannot establish how *s*\* performs when a user specifies a target identity without prior knowledge of a pair’s true divergence. Repeating the comparison at two fixed *a priori* targets, *f*_min_ ∈ {0.90, 0.95}, applied identically across all 54 pairs, coverage depended sharply on whether each pair’s 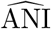 *Ineachof thesedegeneratecases* met the assumption. Notably, coverage was perfect (100%) for pairs at or above target, but falling to 89.5% (*f*_min_ = 0.90) and 78.9% (*f*_min_ = 0.95), below nominal in both cases, for pairs more than 5pp below target, with median realized *ρ* correspondingly inflated (0.185 and 0.357; the latter exceeding the borderline/unreliable boundary; Table 4, Figures S5-S6).

**Table 4.**
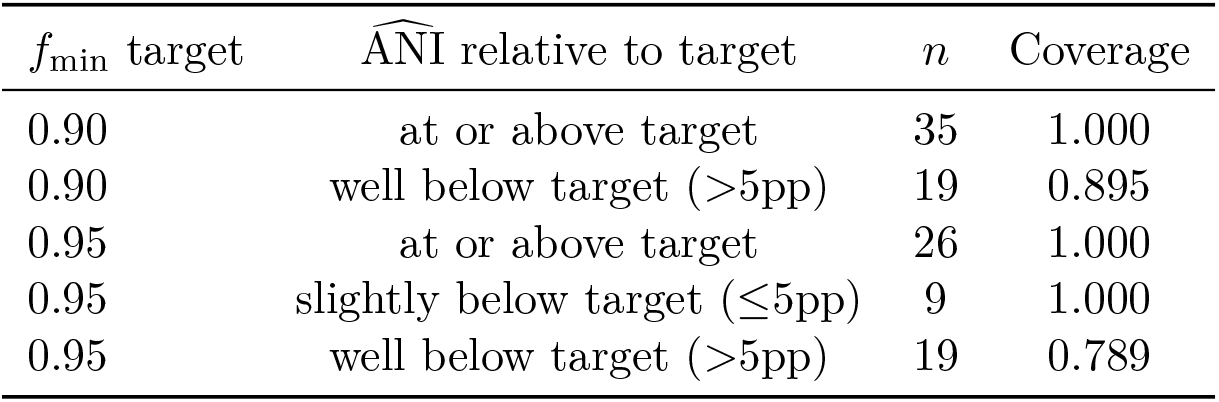
Coverage and precision of *s*\* from fixed *a priori* targets, stratified by whether each pair’s 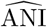met, nearly met, or fell substantially short of the target.

| $f_{\min}$ target | $\widehat{\text{ANI}}$ relative to target | $n$ | Coverage |
| --- | --- | --- | --- |
| 0.90 | at or above target | 35 | 1.000 |
| 0.90 | well below target ($>5\text{pp}$) | 19 | 0.895 |
| 0.95 | at or above target | 26 | 1.000 |
| 0.95 | slightly below target ($\leq 5\text{pp}$) | 9 | 1.000 |
| 0.95 | well below target ($>5\text{pp}$) | 19 | 0.789 |

This establishes that *s*\*’s precision guarantee holds prospectively only when the assumed design target is not violated by a pair’s true divergence, complementing rather than duplicating the retrospective bias result of Section 3.6.

## 4 Discussion

### 4.1 Summary of findings

We set out to answer a precisely stated question: given a genome length *L*, target relative precision *ρ*\*, confidence level 1 − *α*, and *k* selected via Fofanov’s noise-floor criterion, what is the minimum MinHash sketch size *s*\* needed to attain that precision? We show that fixed-size bottom-sketch MinHash admits a direct closed-form answer (Section 3.1), agreeing closely with exact Clopper-Pearson inversion (Section 3.2) and remaining well calibrated on real, overlapping *k*-mers from simulated genomes (Section 3.4). Its main qualification is that the precision calculation itself is reliable conditional on *J*, whereas the biological model used to predict *J* from sequence identity can be substantially biased on real bacterial genomes (Section 3.6).

This closes a specific gap: Hera et al. [2023] derived closed-form confidence intervals for FracMinHash containment but could not obtain the analogous Jaccard interval, since their variable-sketch-size estimator could not be shown asymptotically Normal. Fixed-size bottom-sketch MinHash avoids this obstruction by construction: because both sketches are drawn to a common, predetermined size *s*, the shared-hash count follows a known finite-population distribution (Section 2.1), and the exact Clopper-Pearson interval, and its Wald-based inversion to *s*\*, follow directly.

### 4.2 The formula is reliable, but its inputs are not always

Given a correct value of *J*, the Clopper-Pearson interval retains its coverage and precision guarantee both on idealized draws and on real sketches built from real, overlapping simulated *k*-mers (Section 3.3, Section 3.4). Indeed, coverage remained within 2 percentage points of the idealized reference in 14 of 15 tested cells, the one exception more conservative than nominal. What is not always sound is the exact identity-Jaccard relationship itself, *J*_min_(*k, f*) = *f* ^*k*^*/*(2 − *f* ^*k*^) [Blanca et al., 2022], used to translate a target identity into *J*. This relationship remains accurate to a few percent on simulated sequences (Section 3.4), but breaks down far more severely, and in a taxonomically structured way, on real bacterial genomes (Section 3.6): negligible bias among same-species pairs, growing to +27% at genus level and +77% at family level. This reflects real genome architecture, not the sketching procedure. A plausible explanation is that conserved core genomic regions coexist with more variable, laterally mobile accessory content [Preska Steinberg et al., 2022, Segerman, 2012, Kung et al., 2010], which a single-parameter mutation model necessarily averages over; this is consistent with, though not uniquely diagnostic of, the taxonomic structure observed. Two dataset properties could also contribute in the same direction: 74.2% of assemblies used were contig- or scaffold-level rather than complete (Table S8), and fragmented assemblies may disproportionately undercount accessory content relative to more completely assembled core regions; and 11.3% of assemblies contained unfiltered plasmid sequence. Disentangling these from true core-genome conservation would require completeness- and replicon-stratified analysis on curated, complete genomes, and was beyond the scope of this study. Therefore, we regard core-genome conservation as the best-supported hypothesis rather than a confirmed mechanism. This bias is nonetheless independent evidence, obtained by a different route, of the identity-inflation Hera et al. [2023] report for FracMinHash-based mutation-rate estimation under the same model family, now quantified with an explicit taxonomic gradient.

The practical consequence is favorable: because required sketch size decreases with *J*, and real sub-species pairs have *J*_exact_ systematically *larger* than *J*_theory_, the formula’s error at low identity is conservative rather than a shortfall-consistent with the reliability-tier shift observed at *s*\* versus *s* = 1000 (Section 3.6), though that benchmark alone does not establish tighter calibration, only fewer unreliable classifications. The prospective analysis (Section 3.7) sharpens this: coverage held at 100% for pairs meeting or exceeding an assumed design target, but fell to 78.9-89.5% for pairs whose true identity fell substantially short of it. We characterize *s*\* as a safe, precision-preserving design target whose conservatism depends on the assumed target being met, not a precise *a priori* predictor of what a specific real pair requires below the species level. Together these results separate two sources of uncertainty often conflated in sketch-based comparison: finite-sampling uncertainty, analytically tractable here, and identity-to-similarity modeling uncertainty, which can dominate at larger evolutionary distances.

### 4.3 The staircase structure and its practical implications

That *s*\*(*L*) is piecewise-constant and discontinuous in genome length (Section 3.5) follows directly, and in retrospect elementarily, from Fofanov’s ceiling function. Nonetheless, we are not aware of it being stated explicitly in the sketch-size literature. Its incidence is concentrated rather than uniform: the GTDB interquartile genome-size range (2.17-4.87 Mb) straddles the *k*\* = 13 → 14 boundary at 2.68 Mb directly, a size range in which genome-reduced, host-restricted bacteria are disproportionately represented relative to free-living taxa, making the staircase most relevant precisely for comparisons spanning that biological divide.

A related floor arises at very high identity: 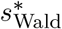 becomes very small (5-12; Section 3.6), where the shared-hash count saturates with probability *J* ^*s*^, 80-90% at the degenerate pairs observed, the modal outcome rather than an edge case. A single fixed floor (e.g. *s* ≥ 100) is insufficient, since the sketch size needed to bound saturation probability grows with *J* and already exceeds such flat floors in this regime. We recommend the explicit floor *s*_sat_(*J, δ*) = ⌈ln *δ/* ln *J* ⌉, applied as 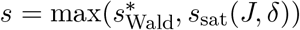 (Supplementary S9). However, avoiding saturation does not by itself make very-high-identity comparisons as precisely characterizable as moderate-divergence ones, since *ρ* remains coarsely discretized at the small sketch sizes this regime calls for.

### 4.4 Limitations

The exact Clopper-Pearson framework applies specifically to fixed-size bottom-sketch Min-Hash, the classical Mash construction [Ondov et al., 2016]. FracMinHash/scaled sketching [Irber, 2020, Irber et al., 2022] is increasingly preferred for cross-size comparisons (e.g. metagenome-to-genome containment), but fixed-size sketching remains the standard for comparing genomes of comparable size, the setting addressed here, and the one underlying FastANI [Jain et al., 2018], still the *de facto* ANI standard, and current fixed-size sketch benchmarking [Zhao et al., 2024]. Our results apply most directly to same-size genome-to-genome comparison (all-pairs clustering, strain delimitation, reference selection), not cross-size metagenomic containment. The same interval machinery does not transfer to FracMinHash, since the shared-hash count no longer has a fixed-*s* finite-population representation there.

The real-genome validation is restricted to bacterial genomes from eight GTDB lineages, with FastANI’s ~ 75-80% resolution floor excluding higher-divergence pairs, and does not extend to archaea, eukaryotes, or viruses. The core/accessory-genome explanation for the observed bias is well supported in direction and magnitude but not independently confirmed via core-genome alignment or annotation. We regard it as the best-supported hypothesis, not an established mechanism. The benchmark itself is a retrospective calibration exercise, 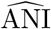used as the plug-in *f*_min_, rather than a full prospective usage test, though Section 3.7 directly addresses the prospective case for two representative targets.

The small-*J* approximations underlying the staircase step-ratio analysis degrade at high minimum identity by an amount equal to *J*_min_(*k, f*) itself (Section 3.1) and should be read as directional only near sequence identity. Finally, all guarantees here are conditional on a correctly specified *k* and target *J*. Indeed, Fofanov’s criterion controls chance *k*-mer collisions but was never intended to correct for the biological non-uniformity of divergence discussed above, and *s*\* is a design requirement conditional on an assumed identity, not a data-independent guarantee for every pair at that identity.

### 4.5 Recommendations and future work

We recommend computing *s*\* directly from the closed form (Section 3.1) using the intended *ρ*\*, *α*, and Fofanov-selected *k*\*(*L*), rather than treating a fixed default such as *s* = 1000 as universally adequate. Where comparisons may span the species boundary or below, *s*\* should be treated as a conservative lower bound rather than a precise target.

Two directions follow. First, empirical correction of the identity-Jaccard relationship for sub-species comparisons, potentially a two-compartment (core/accessory) extension of the simple mutation model, calibrated against the taxonomic gradient reported here, would let *s*\* be tight as well as conservative at high divergence. Second, extending precision-targeted sketch-size (or scale-factor) selection to FracMinHash will require circumventing the asymptotic-normality obstruction identified by Hera et al. [2023], for example via a delta-method or bootstrap-based interval on the Jaccard estimator analogous to their treatment of containment.

## Supporting information

FastANI results for all candidate pairs

Genome accessions used in the real genome benchmark, including the full 96-pair candidate set with shared taxonomic rank

## Data Availability

The code and data that supports the findings of this study are openly available in GitHub at https://github.com/Ebedthan/minhash_s_star.

