## Supplementary material for "How do noise and precision contraints jointly determine the minimum sketch size for fixed-size MinHash Jaccard estimation?"

Anicet E. T. Ebou Dominique K. Koua

#### Contents


**S1. Derivation of the closed-form sketch-size criterion** 2

**S2. Wald approximation versus exact interval inversion, full detail** 3

**S3. Idealized calibration at fixed genome length and identity, full detail** 5

**S4. Formula bias on simulated sequences, by genome length** 6

**S5. Fofanov $k$-band transitions, full detail** 7

**S6. Genome-size distribution and $k$-band boundary proximity** 9

**S7. Genome selection and real genome benchmark composition** 10

**S8. Supplementary figures** 11

**S9. A saturation floor for near-identical comparisons** 13

**S10. Hypergeometric versus binomial sampling** 14

**S11. Equivalence of the bottom-sketch merge construction** 14

**S12. Choice of $L = \max(L_A, L_B)$ for $k$-selection on unequal genome pairs** 15

**S13. Software versions and data availability** 16

#### S1. Derivation of the closed-form sketch-size criterion

For a MinHash sketch of size $s$ and true Jaccard index $J$, the shared-hash count $x$ is modeled as Binomial$(s, J)$-distributed, the standard limiting approximation to the underlying finite-population (hypergeometric) sampling process (Supplementary S10). From this, the derivation of $s^*$ follows.

**Step 1: Wald approximation to the Clopper–Pearson half-width.** The exact Clopper–Pearson interval $[J_{\text{low}}, J_{\text{high}}]$ at confidence level $1-\alpha$ is asymptotically well approximated, for $s$ not too small, by the normal (Wald) interval around $\hat{J} = x/s$:

$$J_{\text{high}} - J_{\text{low}} \;\approx\; 2\, z_{\alpha/2}\, \mathrm{SE}(\hat{J}), \qquad \mathrm{SE}(\hat{J}) = \sqrt{\frac{J(1-J)}{s}}.$$

**Step 2: relative half-width.** By definition, $\rho = (J_{\text{high}} - J_{\text{low}})/(2\hat{J})$. Substituting Step 1 and evaluating $\hat{J} \approx J$ gives

$$\rho \;\approx\; \frac{2\, z_{\alpha/2}\sqrt{J(1-J)/s}}{2J} \;=\; z_{\alpha/2}\sqrt{\frac{1-J}{sJ}}.$$

The half-width is already produced by Step 1 ($z_{\alpha/2}\,\mathrm{SE}(\hat{J})$, not the full width).

**Step 3: inversion for minimum sketch size.** Setting $\rho = \rho^*$ in Step 2 and solving for $s$:

$$\rho^{*2} = z^2_{\alpha/2}\,\frac{1-J}{sJ} \quad\Longrightarrow\quad s^*(J, \rho^*, \alpha) = \frac{z^2_{\alpha/2}(1-J)}{\rho^{*2} J}.$$

**Step 4: composition with the identity–Jaccard and $k$-selection relationships.** Substituting the identity–Jaccard relationship $J = J_{\min}(k, f_{\min}) = f^k_{\min}/(2 - f^k_{\min})$ [1] and Fofanov's $k$-selection criterion $k = k^*(L) = \lceil \log_4(100L) \rceil$ [2] into Step 3 gives the fully joint closed form used to generate all subsequent results in this work:

$$s^*(L, f_{\min}, \rho^*, \alpha) \;=\; \frac{z^2_{\alpha/2}\,(1 - J_{\min}(k^*(L), f_{\min}))}{\rho^{*2}\, J_{\min}(k^*(L), f_{\min})}.$$

**Small-$J$ approximation validity.** The small-$J$ asymptotic approximation used to derive the step-height ratio and advantage-boundary results in the main text, $J_{\min}(k, f) \approx f^k/2$, was checked against the exact expression $J_{\min}(k, f) = f^k/(2 - f^k)$ across the full range of minimum identities and genome lengths used elsewhere in this work ($f_{\min} \in \{0.8, 0.9, 0.95, 0.99, 0.999\}$, $k = k^*(L)$ for $L \in [10^5, 10^{10}]$), given in Table S1. As derived in the main text, the relative error of this approximation is algebraically identical to $J_{\min}(k, f)$ itself. The approximation is tight where $J_{\min}$ is small–under 3.6% relative error at $f_{\min} = 0.8$–and progressively looser at high minimum identity, reaching 96–98% relative error at $f_{\min} = 0.999$, where $J_{\min}$ itself is not small and the approximation is not expected to hold.

Table S1: Relative error of the small-$J$ approximation $J_{\min}(k,f) \approx f^k/2$, across all tested genome lengths and minimum identities.

| $f_{\min}$ | $L$ | $k^*(L)$ | $J_{\min}(k,f)$ (exact) | Relative error (%) |
|---|---|---|---|---|
| 0.999 | $10^5$ | 12 | 0.9764 | 97.6 |
| 0.999 | $10^6$ | 14 | 0.9726 | 97.3 |
| 0.999 | $10^7$ | 15 | 0.9706 | 97.1 |
| 0.999 | $10^8$ | 17 | 0.9668 | 96.7 |
| 0.999 | $10^9$ | 19 | 0.9630 | 96.3 |
| 0.999 | $10^{10}$ | 20 | 0.9611 | 96.1 |
| 0.99 | $10^5$ | 12 | 0.7960 | 79.6 |
| 0.99 | $10^6$ | 14 | 0.7679 | 76.8 |
| 0.99 | $10^7$ | 15 | 0.7545 | 75.4 |
| 0.99 | $10^8$ | 17 | 0.7285 | 72.9 |
| 0.99 | $10^9$ | 19 | 0.7038 | 70.4 |
| 0.99 | $10^{10}$ | 20 | 0.6919 | 69.2 |
| 0.95 | $10^5$ | 12 | 0.3702 | 37.0 |
| 0.95 | $10^6$ | 14 | 0.3225 | 32.2 |
| 0.95 | $10^7$ | 15 | 0.3015 | 30.1 |
| 0.95 | $10^8$ | 17 | 0.2643 | 26.4 |
| 0.95 | $10^9$ | 19 | 0.2326 | 23.3 |
| 0.95 | $10^{10}$ | 20 | 0.2184 | 21.8 |
| 0.90 | $10^5$ | 12 | 0.1644 | 16.4 |
| 0.90 | $10^6$ | 14 | 0.1292 | 12.9 |
| 0.90 | $10^7$ | 15 | 0.1148 | 11.5 |
| 0.90 | $10^8$ | 17 | 0.0910 | 9.1 |
| 0.90 | $10^9$ | 19 | 0.0724 | 7.2 |
| 0.90 | $10^{10}$ | 20 | 0.0647 | 6.5 |
| 0.80 | $10^5$ | 12 | 0.0356 | 3.6 |
| 0.80 | $10^6$ | 14 | 0.0225 | 2.2 |
| 0.80 | $10^7$ | 15 | 0.0179 | 1.8 |
| 0.80 | $10^8$ | 17 | 0.0114 | 1.1 |
| 0.80 | $10^9$ | 19 | 0.0073 | 0.7 |
| 0.80 | $10^{10}$ | 20 | 0.0058 | 0.6 |

### S2. Wald approximation versus exact interval inversion, full detail

Table S2 gives the full breakdown of the ratio $s^*_{\text{exact}}/s^*_{\text{Wald}}$ by $(\rho^*, \alpha)$ combination, including the absolute range of $s^*_{\text{Wald}}$ and $s^*_{\text{exact}}$ spanned within each combination (across the full $J \in [10^{-4}, 0.9]$ grid), so that the practical scale at which any discrepancy occurs can be assessed directly rather than inferred from the ratio alone. The largest relative discrepancies are concentrated at the smallest absolute sketch sizes in the grid (single digits to low tens, arising at the largest tested $J$ under the loosest confidence settings) and are correspondingly of limited practical consequence; agreement is uniformly close at the sketch sizes most relevant to genome comparison in practice ($s^*$ in the hundreds to low millions, arising at small-to-moderate $J$).

Across the full evaluated grid, 341 of 360 cells were valid after excluding 19 combinations infeasible within the $s_{\max} = 5 \times 10^6$ search bound described in the main text Methods. Agreement between $s^*_{\text{Wald}}$ and $s^*_{\text{exact}}$ was tightest at the tighter, more commonly used precision targets: at $\rho^* = 0.05$ the median relative error ranged from 0.8% ($\alpha = 0.01$) to 1.9% ($\alpha = 0.10$), and at $\rho^* = 0.10$–the target used as the default "reliable" threshold throughout this work–from 1.6% to 3.8% (Table S2). Error increased systematically with looser $\rho^*$ and looser (larger) $\alpha$, consistent

Table S2: Ratio of exact to Wald-approximated minimum sketch size, with the absolute range of $s^*_{\text{Wald}}$ and $s^*_{\text{exact}}$ spanned across the $J \in [10^{-4}, 0.9]$ grid (40 log-spaced points) for each $(\rho^*, \alpha)$ combination. RE: relative error.

| $\rho^*$ | $\alpha$ | $n$ cells | $s^*_{\text{Wald}}$ range | | $s^*_{\text{exact}}$ range | | median RE (%) | max RE (%) | max ratio |
|---|---|---|---|---|---|---|---|---|---|
| | | | min | max | min | max | | | |
| 0.05 | 0.01 | 32 | 295 | 4,097,192 | 313 | 4,128,489 | 0.79 | 6.14 | 1.061 |
| 0.05 | 0.05 | 35 | 171 | 4,780,380 | 191 | 4,843,898 | 1.35 | 11.87 | 1.119 |
| 0.05 | 0.10 | 36 | 120 | 4,252,464 | 141 | 4,333,266 | 1.90 | 17.26 | 1.173 |
| 0.10 | 0.01 | 38 | 74 | 4,158,945 | 82 | 4,222,367 | 1.59 | 11.23 | 1.112 |
| 0.10 | 0.05 | 40 | 43 | 3,841,075 | 52 | 3,945,001 | 2.76 | 21.83 | 1.218 |
| 0.10 | 0.10 | 40 | 30 | 2,705,273 | 40 | 2,805,001 | 3.81 | 33.06 | 1.331 |
| 0.30 | 0.01 | 40 | 8 | 737,137 | 11 | 775,000 | 5.15 | 34.29 | 1.343 |
| 0.30 | 0.05 | 40 | 5 | 426,786 | 8 | 465,001 | 9.01 | 68.69 | 1.687 |
| 0.30 | 0.10 | 40 | 3 | 300,586 | 7 | 335,000 | 11.57 | 109.57 | 2.096 |

with the Wald approximation's known degradation as the effective sample size implied by the target shrinks.

The single cell in which $s^*_{\text{exact}}$ fell below $s^*_{\text{Wald}}$ occurred at $J = 0.7126$, $\rho^* = 0.30$, $\alpha = 0.01$, where $s^*_{\text{exact}} = 29$ fell fractionally below $s^*_{\text{Wald}} \approx 29.7$ (ratio 0.975). This cell was one of 37 (10.9% of valid cells) in which $s^*_{\text{exact}}$ and $s^*_{\text{stable}}$–the smallest sketch size beyond which the target is met for all larger $s$–differed. This confirms the non-monotonicity of $\rho_{\text{exact}}(s, J, \alpha)$ noted in the main text: because $x = \text{round}(Js)$ introduces integer rounding, the exact relative half-width is not guaranteed to decrease monotonically in $s$, and at this cell the criterion was transiently satisfied at $s = 29$, violated again at $s = 30$, and held permanently only from $s = 31$ onward ($s^*_{\text{stable}} = 31$, itself exceeding $s^*_{\text{Wald}}$ as expected). Across all 37 cells where the two quantities differed, the gap ranged from 2 to 380 integer steps (median 3), concentrated at small $J$ and loose $(\rho^*, \alpha)$ settings; in the remaining 89.1% of valid cells, $s^*_{\text{exact}}$ and $s^*_{\text{stable}}$ coincided exactly. This confirms that the isolated small-$s$ discrepancy reflects genuine, if rare, integer-rounding non-monotonicity in the exact criterion rather than any deficiency in the Wald approximation itself, which otherwise held its conservative margin throughout the grid.

#### S3. Idealized calibration at fixed genome length and identity, full detail

Table S3 shows coverage and realized $\rho$ across the $(\rho^*, \alpha)$ sweep at $L = 1$ Mb, $f_{\min} = 0.9$ ($J = 0.1292$). Across the full $(L, f_{\min})$ grid at $\rho^* = 0.10$, $\alpha = 0.05$ (150 cells), coverage ranged from 0.9475 to 0.9984 (median 0.9553) and median realized $\rho$ across cells was 0.1017 (Figures S1–S2).

Table S3: Empirical calibration of $s^*_{\text{Wald}}$ under the idealized Binomial($s^*$, $J$) model at $L = 1$ Mb, $f_{\min} = 0.9$ ($J = 0.1292$).

| $\rho^*$ | $\alpha$ | $s^*$ | Coverage (target $1-\alpha$) | Median realized $\rho$ |
|---|---|---|---|---|
| 0.05 | 0.01 | 17,895 | 0.991 (0.99) | 0.0502 |
| 0.05 | 0.05 | 10,361 | 0.951 (0.95) | 0.0504 |
| 0.05 | 0.10 | 7,297 | 0.906 (0.90) | 0.0505 |
| 0.10 | 0.01 | 4,474 | 0.991 (0.99) | 0.1016 |
| 0.10 | 0.05 | 2,591 | 0.952 (0.95) | 0.1016 |
| 0.10 | 0.10 | 1,825 | 0.906 (0.90) | 0.1020 |
| 0.30 | 0.01 | 498 | 0.989 (0.99) | 0.3079 |
| 0.30 | 0.05 | 288 | 0.965 (0.95) | 0.3138 |
| 0.30 | 0.10 | 203 | 0.905 (0.90) | 0.3198 |

Despite the small, consistent median overshoot of realized $\rho$ above $\rho^*$ reported in the main text, the proportion of individual sketching replicates achieving $\rho \leq \rho^*$ was itself below 50% in every tested cell (e.g. 29.98% at $\rho^* = 0.10$, $\alpha = 0.05$, $L = 1$ Mb, $f_{\min} = 0.9$), a property of the discreteness of $\rho$ as a function of the integer-valued shared-hash count $x$, rather than an indication of miscalibration. Direct inspection of the empirical quantiles of realized $\rho$ at this representative cell confirmed this: the 20th, 30th, and 50th percentiles were 0.0990, 0.1000, and 0.1016 respectively, showing that the empirical distribution function of $\rho$ rises steeply through the target value, consistent with $\rho(x)$ being a smooth, nearly linear, monotonically decreasing function of $x$ near the mode of a moderately large binomial count, such that a small shift in the median relative to $\rho^*$ corresponds to a much larger shift in the proportion of replicates falling below it. No cell showed a directional inconsistency between the sign of the median-$\rho$ overshoot and the sign of the corresponding proportion-below-target deviation from 50% (0 of 150 cells flagged).

#### S4. Formula bias on simulated sequences, by genome length

Table S4 gives the $J_{\text{theory}}$ vs. $J_{\text{exact}}$ bias on simulated sequences broken out by genome length rather than summarized as a median, showing that the bias trend with minimum identity is consistent across the three tested genome lengths (Figure S3).

Table S4: Relative bias of the fixed formula prediction $J_{\text{theory}}$ against the realized $J_{\text{exact}}$ on simulated sequences, by genome length and minimum identity.

| $L$ | $f_{\text{min}}$ | $J$ formula bias (%) |
|---|---|---|
| $10^4$ | 0.999 | 0.02 |
| $10^4$ | 0.99 | 0.10 |
| $10^4$ | 0.95 | 0.79 |
| $10^4$ | 0.90 | 2.14 |
| $10^4$ | 0.80 | 7.80 |
| $10^5$ | 0.999 | 0.01 |
| $10^5$ | 0.99 | 0.08 |
| $10^5$ | 0.95 | 0.52 |
| $10^5$ | 0.90 | 1.54 |
| $10^5$ | 0.80 | 8.07 |
| $10^6$ | 0.999 | 0.01 |
| $10^6$ | 0.99 | 0.06 |
| $10^6$ | 0.95 | 0.39 |
| $10^6$ | 0.90 | 1.27 |
| $10^6$ | 0.80 | 8.13 |

#### S5. Fofanov $k$-band transitions, full detail

Table S5 gives all 40 detected $k$-band transitions underlying the main-text step-height ratio table and staircase figure: the predicted crossing point $L = 4^k/100$, the grid bracket confirming it, and the observed versus predicted step-height ratio for every transition. The predicted crossing point fell inside the grid's bracketing interval for all 40 transitions (maximum bracket width 0.58% of $L_{\text{predicted}}$).

Table S5: All detected Fofanov $k$-band transitions: predicted crossing point $L = 4^k/100$, grid bracket confirming it falls within the tested resolution, and observed vs. predicted $(1/f_{\min})$ step-height ratio.

| $f_{\min}$ | $k \to k{+}1$ | $L_{\text{predicted}}$ | Bracket width (%) | Obs. ratio | Pred. ratio | Rel. error (%) |
|---|---|---|---|---|---|---|
| 0.8 | 12→13 | 167,772 | 0.57 | 1.268 | 1.250 | 1.48 |
| 0.8 | 13→14 | 671,089 | 0.58 | 1.265 | 1.250 | 1.16 |
| 0.8 | 14→15 | 2,684,355 | 0.58 | 1.262 | 1.250 | 0.92 |
| 0.8 | 15→16 | 10,737,418 | 0.57 | 1.259 | 1.250 | 0.73 |
| 0.8 | 16→17 | 42,949,673 | 0.58 | 1.257 | 1.250 | 0.58 |
| 0.8 | 17→18 | 171,798,692 | 0.58 | 1.256 | 1.250 | 0.46 |
| 0.8 | 18→19 | 687,194,767 | 0.58 | 1.255 | 1.250 | 0.37 |
| 0.8 | 19→20 | 2,748,779,069 | 0.58 | 1.254 | 1.250 | 0.29 |
| 0.9 | 12→13 | 167,772 | 0.57 | 1.155 | 1.111 | 3.94 |
| 0.9 | 13→14 | 671,089 | 0.58 | 1.149 | 1.111 | 3.41 |
| 0.9 | 14→15 | 2,684,355 | 0.58 | 1.144 | 1.111 | 2.97 |
| 0.9 | 15→16 | 10,737,418 | 0.57 | 1.140 | 1.111 | 2.59 |
| 0.9 | 16→17 | 42,949,673 | 0.58 | 1.136 | 1.111 | 2.27 |
| 0.9 | 17→18 | 171,798,692 | 0.58 | 1.133 | 1.111 | 2.00 |
| 0.9 | 18→19 | 687,194,767 | 0.58 | 1.131 | 1.111 | 1.77 |
| 0.9 | 19→20 | 2,748,779,069 | 0.58 | 1.128 | 1.111 | 1.56 |
| 0.95 | 12→13 | 167,772 | 0.57 | 1.115 | 1.053 | 5.88 |
| 0.95 | 13→14 | 671,089 | 0.58 | 1.108 | 1.053 | 5.27 |
| 0.95 | 14→15 | 2,684,355 | 0.58 | 1.103 | 1.053 | 4.76 |
| 0.95 | 15→16 | 10,737,418 | 0.57 | 1.098 | 1.053 | 4.32 |
| 0.95 | 16→17 | 42,949,673 | 0.58 | 1.094 | 1.053 | 3.93 |
| 0.95 | 17→18 | 171,798,692 | 0.58 | 1.090 | 1.053 | 3.59 |
| 0.95 | 18→19 | 687,194,767 | 0.58 | 1.087 | 1.053 | 3.29 |
| 0.95 | 19→20 | 2,748,779,069 | 0.58 | 1.085 | 1.053 | 3.03 |
| 0.99 | 12→13 | 167,772 | 0.57 | 1.089 | 1.010 | 7.80 |
| 0.99 | 13→14 | 671,089 | 0.58 | 1.082 | 1.010 | 7.16 |
| 0.99 | 14→15 | 2,684,355 | 0.58 | 1.077 | 1.010 | 6.62 |
| 0.99 | 15→16 | 10,737,418 | 0.57 | 1.072 | 1.010 | 6.15 |
| 0.99 | 16→17 | 42,949,673 | 0.58 | 1.068 | 1.010 | 5.73 |
| 0.99 | 17→18 | 171,798,692 | 0.58 | 1.064 | 1.010 | 5.37 |
| 0.99 | 18→19 | 687,194,767 | 0.58 | 1.061 | 1.010 | 5.04 |
| 0.99 | 19→20 | 2,748,779,069 | 0.58 | 1.058 | 1.010 | 4.75 |
| 0.999 | 12→13 | 167,772 | 0.57 | 1.084 | 1.001 | 8.28 |
| 0.999 | 13→14 | 671,089 | 0.58 | 1.077 | 1.001 | 7.64 |
| 0.999 | 14→15 | 2,684,355 | 0.58 | 1.072 | 1.001 | 7.09 |
| 0.999 | 15→16 | 10,737,418 | 0.57 | 1.067 | 1.001 | 6.61 |
| 0.999 | 16→17 | 42,949,673 | 0.58 | 1.063 | 1.001 | 6.20 |
| 0.999 | 17→18 | 171,798,692 | 0.58 | 1.059 | 1.001 | 5.83 |
| 0.999 | 18→19 | 687,194,767 | 0.58 | 1.056 | 1.001 | 5.50 |
| 0.999 | 19→20 | 2,748,779,069 | 0.58 | 1.053 | 1.001 | 5.21 |

#### S6. Genome-size distribution and $k$-band boundary proximity

To assess how frequently real genomes fall near a Fofanov $k$-band boundary, we obtained genome sizes for the complete set of genomes in GTDB release 232 (901,341 genomes: 878,998 bacteria and 22,343 archaea) from GTDB's bulk metadata tables (`bac120_metadata_r232.tsv`, `ar53_metadata_r232.tsv`) [3]. For each genome, $k^*(L)$ was computed via Fofanov's criterion from its reported genome size, and the genome was classified as falling within a given tolerance (5%, 10%, or 20%) of a $k$-band boundary $L = 4^k/100$ if its genome size fell within that relative distance of any boundary in the range spanned by GTDB genome sizes. This was repeated separately for the full genome set and for the subset of 199,923 GTDB species representative genomes, to check whether the resulting proportions were sensitive to the uneven sequencing effort across taxa (e.g. the disproportionate representation of frequently sequenced pathogens such as *Escherichia coli*) rather than reflecting the underlying genome-size distribution itself.

Across the full census, 99.06% of genomes fell within a single, wide $k^*$ band; 7.6% fell within 5% of a boundary and 15.5% within 10%, essentially unchanged when restricted to species representatives alone (7.8% and 15.5%, respectively), indicating this is a property of the genome-size distribution itself rather than an artifact of uneven sequencing effort.

*Note on sampling precision.* A simple-random-sample estimate of these proportions at $\pm 0.05$ percentage-point precision (95% CI) would require approximately 730,000 of GTDB's $\sim$900,000 genomes–81% of the population–under worst-case variance assumptions. At this precision target, sampling offers no meaningful efficiency gain over a full census; we therefore used the complete genome-size census directly rather than an API-sampled subset.

#### S7. Genome selection and real genome benchmark composition

Table S6 lists the eight anchor bacterial species used for genome sampling. For each anchor, the GTDB species representative genome was retrieved and 12 additional genomes were sampled (three each at species, genus, family, and order rank), yielding 96 candidate pairs in total; order-rank pairs and pairs falling below FastANI's effective resolution floor were excluded, leaving the 54 pairs analyzed in the real genome benchmark section of the main text. The full list of 96 candidate pairs, including accessions and shared taxonomic rank, is provided as a supplementary data file (`genome_selection.tsv`).

Table S6: Anchor bacterial species and their GTDB species representative genome accession.

| Anchor species | Representative accession |
|---|---|
| *Escherichia coli* | GCF_003697165.2 |
| *Staphylococcus aureus* | GCF_001027105.1 |
| *Pseudomonas aeruginosa* | GCF_001457615.1 |
| *Salmonella enterica* | GCF_000006945.2 |
| *Bacillus subtilis* | GCF_000009045.1 |
| *Klebsiella pneumoniae* | GCF_000742135.1 |
| *Mycobacterium tuberculosis* | GCF_000195955.2 |
| *Streptococcus pneumoniae* | GCF_001457635.1 |

Table S7 gives the exact per-rank bias values underlying the main-text discussion of taxonomic structure in the $J_{\text{theory}}$ vs. $J_{\text{exact}}$ bias, confirmed directly against the 54 retained pairs joined with their shared taxonomic rank.

Table S7: Median $J_{\text{theory}}$ vs. $J_{\text{exact}}$ bias on the 54 real genome pairs, by shared taxonomic rank with the anchor genome.

| Shared rank | $n$ pairs | Median $\widehat{\text{ANI}}$ | Median $J$ bias (%) |
|---|---|---|---|
| Species | 24 | 0.987 | −1.37 |
| Genus | 17 | 0.910 | +27.11 |
| Family | 13 | 0.809 | +77.09 |

Table S8 gives the assembly-level composition of the 62 distinct genome assemblies used in the 54 retained pairs, retrieved via `datasets summary genome accession`.

Table S8: Assembly-level composition of the 62 genome assemblies used in the real genome benchmark.

| Assembly level | $n$ | % |
|---|---|---|
| Chromosome | 1 | 1.6 |
| Complete Genome | 15 | 24.2 |
| Contig | 33 | 53.2 |
| Scaffold | 13 | 21.0 |
| **Total** | 62 | 100.0 |

#### S8. Supplementary figures

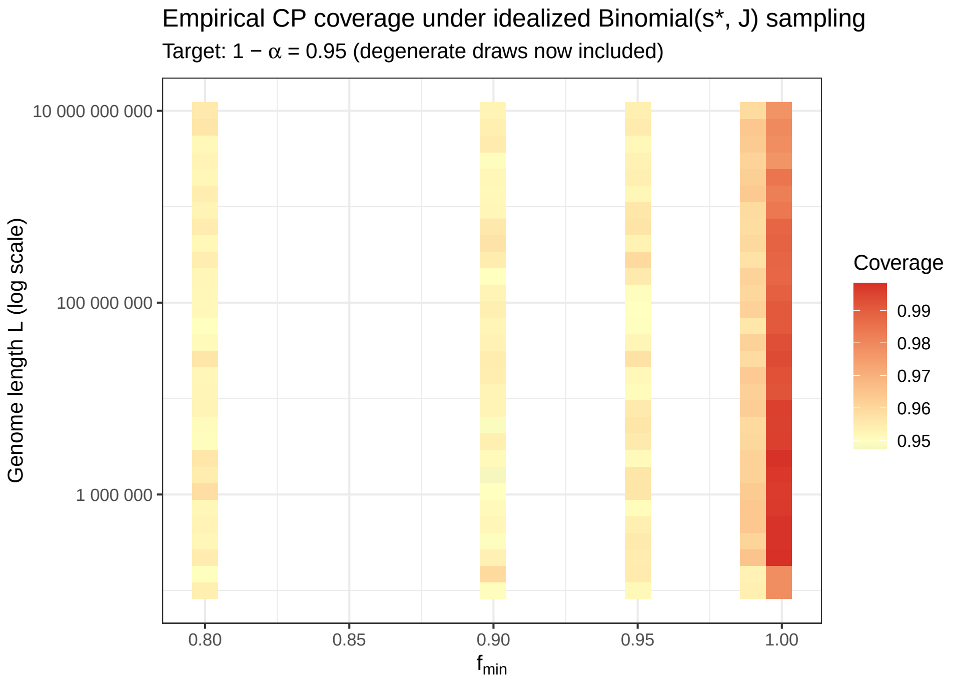


Figure S1: Empirical Clopper–Pearson coverage under the idealized Binomial($s^*$, $J$) model, across the full $(L, f_{\min})$ grid at $\rho^* = 0.10$, $\alpha = 0.05$.

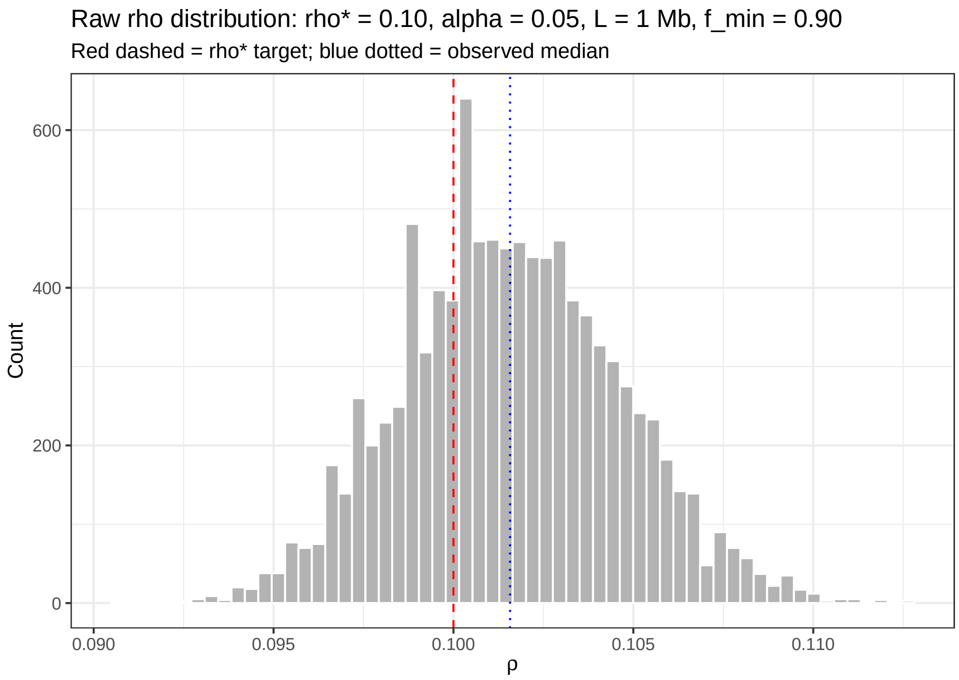


Figure S2: Empirical distribution of realized $\rho$ at the representative cell ($\rho^* = 0.10$, $\alpha = 0.05$, $L = 1$ Mb, $f_{\min} = 0.9$), illustrating the discreteness effect described in Supplementary S3: the median realized $\rho$ sits close to but slightly above the target $\rho^*$, while the proportion of individual replicates falling at or below $\rho^*$ is well below 50% due to the steepness of the empirical distribution function near the target.

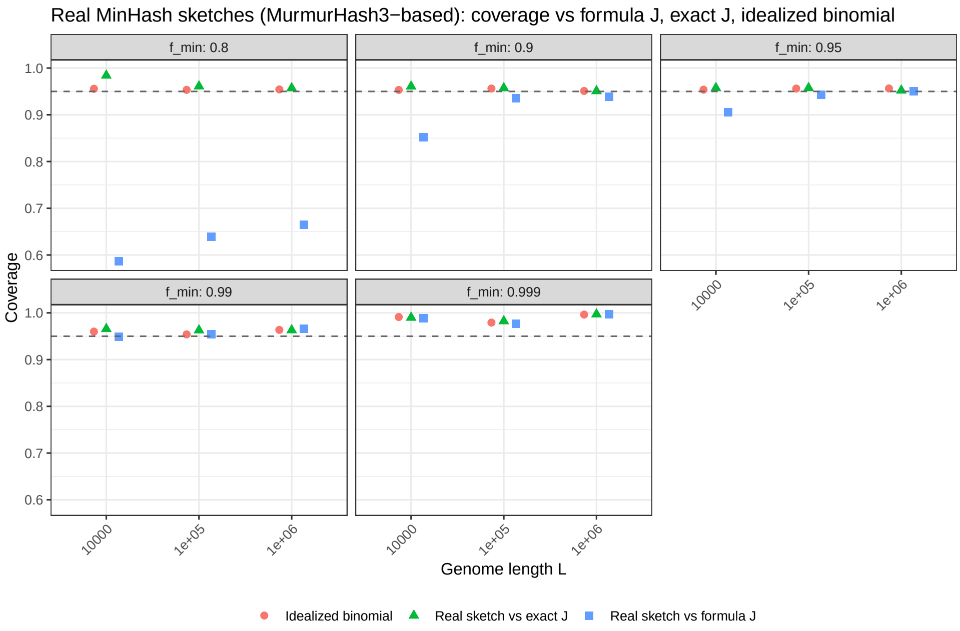


Figure S3: Coverage of the Clopper–Pearson interval against the fixed formula prediction $J_{\text{theory}}$, each replicate's exact realized Jaccard index $J_{\text{exact}}$, and the idealized Binomial($s^*$, $J_{\text{theory}}$) reference, on simulated sequences across all tested genome lengths and minimum identities.

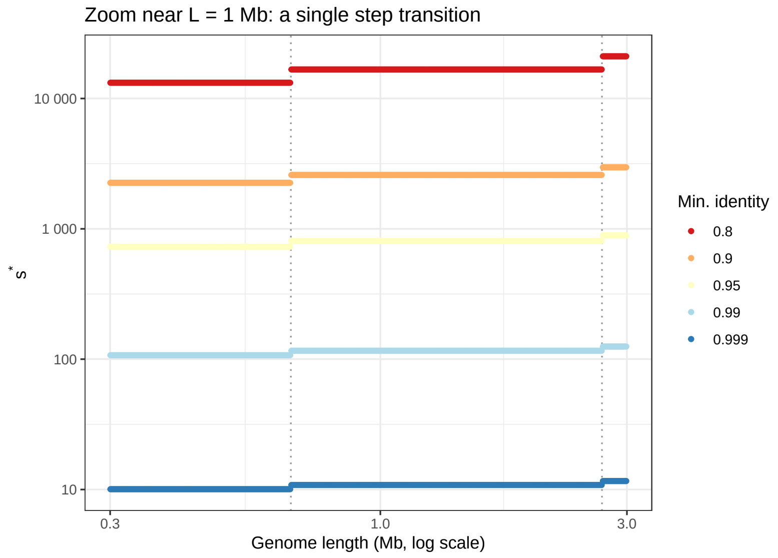


Figure S4: Zoomed view of the staircase structure of $s^*(L)$ near $L = 1$ Mb, showing a single step transition in detail.

#### S9. A saturation floor for near-identical comparisons

At very high minimum identity ($f_{\min} \gtrsim 0.999$), $J_{\min}(k, f_{\min})$ approaches 1 and the closed-form $s^*_{\text{Wald}}$ correspondingly becomes very small (5–12 in the main text's real genome benchmark). This reflects the fact that, under the model, few samples are needed to achieve a small *relative* half-width when $J$ itself is close to 1. This creates a distinct failure mode from the identity–Jaccard bias discussed elsewhere in this work: under the idealized Binomial$(s, J)$ model, the shared-hash count saturates ($x = s$, giving $\hat{J} = 1$ and an undefined $\rho$) with probability $J^s$, which is substantial precisely when $s$ is small and $J$ is close to 1–exactly the regime that $s^*_{\text{Wald}}$ selects at very high identity. At the four degenerate pairs observed in the main text's real genome benchmark ($J_{\text{exact}} = 0.980$–$0.987$, $s^* = 5$–$12$), this saturation probability was 80–90%, meaning saturation is the modal outcome at the formula's own recommended sketch size in this regime, not a rare tail event.

A simple, uniform floor on $s^*$ is not by itself sufficient to control this: because $J^s$ decays slowly in $s$ near $J = 1$, the sketch size needed to keep saturation probability below a chosen tolerance $\delta$ grows with $J$ and is generally larger than commonly proposed fixed floors. Solving $J^s \leq \delta$ gives an explicit, $J$-dependent saturation floor,

$$s_{\text{sat}}(J, \delta) = \left\lceil \frac{\ln \delta}{\ln J} \right\rceil,$$

which at $\delta = 0.05$ requires $s \geq 99$ at $J = 0.97$, $s \geq 149$ at $J = 0.98$, and $s \geq 299$ at $J = 0.99$–already exceeding a flat floor of $s = 100$ for most of this range, and growing further as $\delta$ is tightened (e.g. $s \geq 228$ at $J = 0.98$, $\delta = 0.01$). We therefore recommend that, rather than a single fixed floor, practitioners apply $s = \max(s^*_{\text{Wald}}(J, \rho^*, \alpha),\ s_{\text{sat}}(J, \delta))$ for a chosen saturation tolerance $\delta$ (e.g. $\delta = 0.05$), so that the precision-driven design formula and an explicit saturation-avoidance constraint are both satisfied. We note that this floor does not solve the underlying interpretability problem when saturation does occur: even with $x < s$, $\hat{J}$ near 1 combined with small $s$ still yields a wide, discretely-stepped $\rho$ (Supplementary S3), so very high identity comparisons remain intrinsically harder to characterize precisely than moderate-divergence ones, regardless of sketch size.

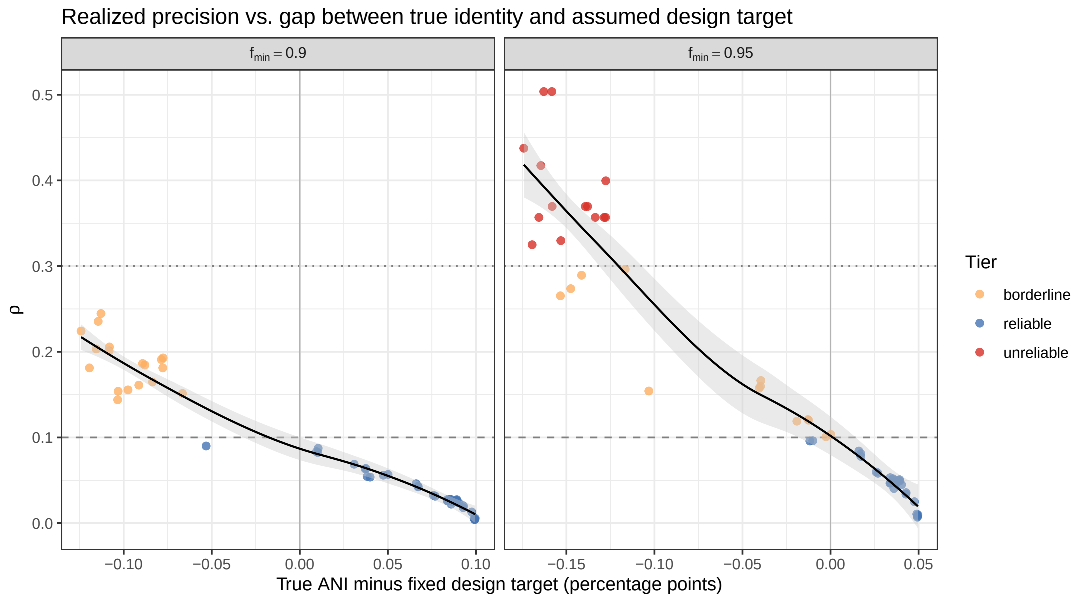


Figure S5: Realized relative half-width $\rho$ as a continuous function of the gap between each pair's true ANI (FastANI) and the fixed *a priori* design target $f_{\min}$, for both tested targets ($f_{\min} = 0.90$, $f_{\min} = 0.95$). Points are coloured by reliability tier; dashed and dotted horizontal lines mark the reliable ($\rho \leq 0.10$) and borderline ($\rho \leq 0.30$) boundaries used throughout this work. The black curve is a loess smooth with 95% pointwise confidence ribbon, shown for visual guidance only given the modest sample size ($n = 54$ pairs per target).

### S10. Hypergeometric versus binomial sampling

Bottom-$s$ sketches sample $s$ elements without replacement from the finite union $H(A) \cup H(B)$ of size $N$, of which $K = |A \cap B|$ are shared; the shared-hash count is therefore exactly $x \sim \text{Hypergeometric}(N, K, s)$ [1], not binomial. We model $x$ as $\text{Binomial}(s, J)$ with $J = K/N$ throughout this work, the standard limiting approximation as $N \to \infty$ at fixed $J$. Because hypergeometric sampling has variance $sJ(1-J)(N-s)/(N-1)$, strictly smaller than the binomial variance $sJ(1-J)$ at the same $N, K, s$ for any finite $N > s$, this approximation makes the resulting Clopper–Pearson interval conservative (wider than strictly necessary) rather than anti-conservative. The approximation is negligible in this work given that $N$ (genome-scale, typically $10^4$–$10^7$ $k$-mers) vastly exceeds $s$ throughout.

### S11. Equivalence of the bottom-sketch merge construction

Any element of $\text{bottom}_s(H(A) \cup H(B))$ must, by definition of rank, already lie in $H_A$ or $H_B$ (if it lies in $H(A)$, it has fewer than $s$ elements of $H(A) \cup H(B)$–and hence of $H(A)$–below it, so it lies in $H_A = \text{bottom}_s(H(A))$; symmetrically for $H(B)$). Thus $\text{bottom}_s(H(A) \cup H(B)) \subseteq H_A \cup H_B$, and taking $\text{bottom}_s(H_A \cup H_B)$ recovers it exactly. This construction is therefore exact, not an approximation, and does not correspond to the alternative, non-equivalent procedure of independently sketching each genome and directly intersecting the two fixed-size sketches without re-sketching their union.

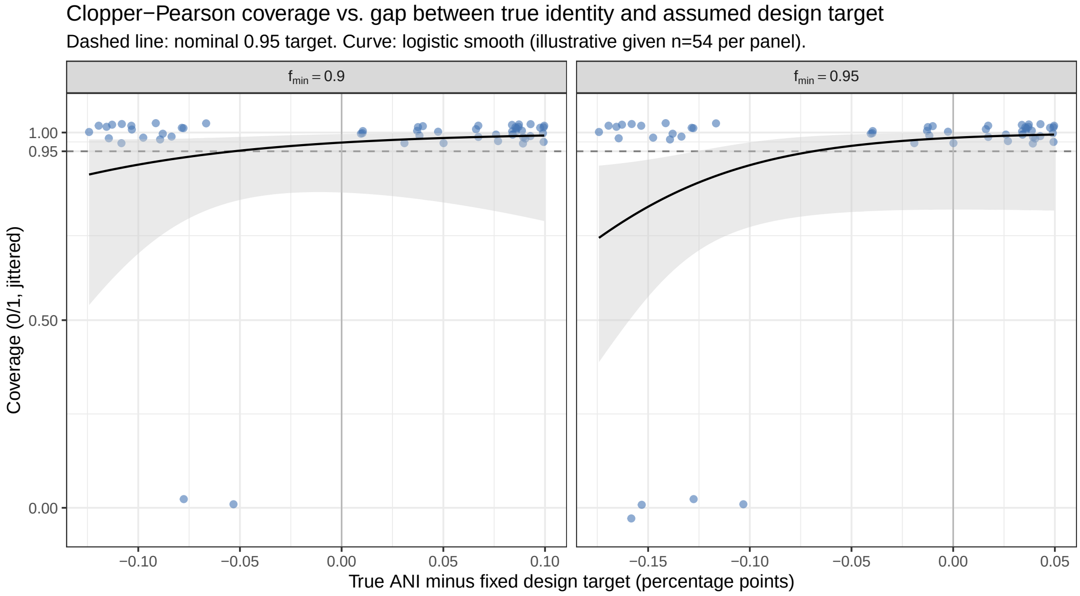


Figure S6: Clopper–Pearson coverage (binary, jittered for visibility) as a function of the same ANI gap as Figure S5, with a logistic regression smooth and 95% confidence ribbon shown for visual guidance only. Coverage is high and stable for pairs at or above the assumed design target and declines for pairs whose true identity falls substantially below it, consistent with the binned summary in the main text.

#### S12. Choice of $L = \max(L_A, L_B)$ for $k$-selection on unequal genome pairs

Fofanov's criterion is derived for two genomes of equal length $L$. Its natural two-length generalization bounds the expected number of chance-shared $k$-mers by $L_A L_B/4^k$, giving $k \geq \log_4(100\, L_A L_B)$. Since $L_A L_B \leq \max(L_A, L_B)^2$, using $L = \max(L_A, L_B)$ does not guarantee $k^*(\max(L_A, L_B)) \geq \log_4(100\, L_A L_B)$ in general, but the two selections coincide when $L_A \approx L_B$, as was the case for all pairs in this dataset: $k^*(\max(L_A, L_B))$ and $k^*(L_A)$ agreed exactly for all 54 retained pairs.

#### S13. Software versions and data availability

Genome selection was performed against GTDB release 232, queried via `xgt` v1.1; genome sequences were downloaded via the NCBI `datasets` command-line tool v18.34.0. Average nucleotide identity was estimated with FastANI v1.33. All analyses were implemented in R v4.6.1, using `qbeta`/`qnorm` for exact and asymptotic quantile computation, `digest` for MurmurHash3-based hashing, `Biostrings` for genome sequence handling, `furrr` v0.4.0/`future` v1.75.0 for parallel Monte Carlo replication, and `ggplot2` v4.0.3 for visualization.

All analysis code, raw output files (including all files underlying the tables and figures in this document), and the reference implementation of $s^*(L, f_{\min}, \rho^*, \alpha)$ are available at [repository URL]. Genome accessions used in the real genome benchmark, including the full 96-pair candidate set with shared taxonomic rank (`genome_selection.tsv`) and FastANI results for all candidate pairs (`fastani_results.tsv`), are provided as supplementary data files at the same location.
